# Visuo-motor rotation influences representational acuity but not space representation

**DOI:** 10.1101/2020.04.06.027508

**Authors:** C Michel, S Amoura, O White

## Abstract

Prism adaptation is a well-known experimental procedure to study sensorimotor plasticity. It has been shown that following prism exposure, after-effects are not only restricted to the sensorimotor level but extend as well to spatial cognition. In the present study, we used a visuo-motor rotation task which approaches the perturbations induced by prism exposure. We induced either leftward or rightward 15-degree rotations and we presented the perturbation either abruptly (from one trial to the next) or gradually (over a 34-trial transition). First, we found that none of the conditions produced cognitive after-effects in perceptive line bisection task. This result has a strong methodological impact for prospective investigations focusing on sensorimotor plasticity while sparing space cognition; it is particularly relevant when investigating sensorimotor plasticity in patients with specific representational feature to preserve from aggravation. Second, another interesting result was the increase of the sensitivity with which we discriminate the center of the line, that we propose to call representational acuity. It improved following the perturbation more particularly after gradual exposure and persisted for some time after the sensorimotor adaptation. These innovative results are discussed in terms of sensorimotor processes underpinning the transfer of visuomotor plasticity to spatial cognition.

## Introduction

Humans tend to preferentially focus their attention to the left side of space. This phenomenon occurs because the right hemisphere is dominant in visuospatial processes (Fink, Marshall, Weiss, & Zilles, 2001). This leftward spatial bias has been referred to as pseudoneglect (Bowers & Heilman, 1980) by analogy to the performance of right-hemisphere impaired patients with left unilateral spatial neglect who show spatial larger biases towards the right side of the space (T. C. Nijboer, Kollen, & Kwakkel, 2013; Rode, Pagliari, Huchon, Rossetti, & Pisella, 2017). Pseudoneglect is particularly well documented for space representation (mental image of the space mapped across the brain) (e.g. McCourt & Jewell, 1999) that is classically assessed and well quantified with a line bisection task. In that test, the estimation of the center of lines is usually characterized by a leftward bias corresponding to a mental overrepresentation of the left part of the space and a mental underpresentation of the right part of the space (Brooks, Della Sala, & Darling, 2014; McCourt & Jewell, 1999). Pseudoneglect generalizes beyond direct visuospatial processes. For instance, it occurs as well on the representation of stimuli with spatial association, i.e. stimuli that are not intrinsically spatial but considered as activating a mental spatial representation with a left to right continuum. Such examples are the observation of a bias toward small numbers in a series of numbers (Loftus, Nicholls, Mattingley, Chapman, & Bradshaw, 2009; Longo & Lourenco, 2007), towards earlier alphabetic letters (Zorzi, Priftis, Meneghello, Marenzi, & Umilta, 2006) or even toward low auditory frequencies ((Michel, Bonnet, Podor, Bard, & Poulin-Charronnat, 2019).

Researchers attempted to modulate this bias with experimental paradigms. Interestingly, sensorimotor plasticity induced by a short period of visuo-manual pointing while wearing goggles that shift the visual field to the left, reverses pseudoneglect behavior into neglect-like behavior (Michel, 2006; 2016 for reviews). In that case, there is a mental overrepresentation of the right part of the space and a mental underrepresentation of the left part of the space (Colent, Pisella, Bernieri, Rode, & Rossetti, 2000; Fortis, Goedert, & Barrett, 2011; Michel, Pisella, et al., 2003; Michel, Rossetti, Rode, & Tilikete, 2003; T. Nijboer, Vree, Dijkerman, & Van der Stigchel, 2010; Schintu et al., 2014; Striemer & Danckert, 2010). Neglect simulation, expressed by a response bias toward the right space, was not only described in peripersonal space representation, but also occurred in extrapersonal (Berberovic & Mattingley, 2003; Michel, Vernet, Courtine, Ballay, & Pozzo, 2008) and bodily space representations (Michel, Rossetti, et al., 2003). Prism adaptation also produces biases toward features with rightward spatial association by shifting the response biases toward large numbers (Loftus, Nicholls, Mattingley, & Bradshaw, 2008), later letters (Nicholls, Kamer, & Loftus, 2008) or higher auditory frequencies (Michel et al., 2019). Furthermore, prism adaptation to a rightward optical deviation is successfully used in neglect rehabilitation (e.g. Jacquin-Courtois et al., 2013; Rossetti et al., 1998).

While prism adaptation induces cognitive after-effects on space representation, a natural following question is whether other kinds of perturbations share the same properties. The fundamental question is to identify the relevant signal involved in sensorimotor plasticity that is responsible for representational after-effects. A recent experiment made a first step toward that direction by assessing whether representational after-effects occurred following adaptation to a novel dynamic environment producing proprioceptive changes. Indeed, even if adaptation to a novel dynamic environment produced change in egocentric reachability judgment suggesting the updating of the arm’s internal model of limb dynamics (Leclere, Sarlegna, Coello, & Bourdin, 2019), it failed to show representational after-effects assessed with allocentric perceptual line bisection task (Michel, Bonnetain, Amoura, & White, 2018). These results in line bisection suggest that the main difference between prism and force field adaptation may lie in the relative contribution of the recalibration and realignment processes. Calibration (fast process) allows to reduce the error by using feedback (proprioceptive and/or visual) and contributes little to sensorimotor after-effects. In parallel, realignment (slow process) brings the different reference frames back in congruence. It develops gradually and is mainly responsible for sensorimotor after-effects (O’Shea et al., 2014). In prism adaptation, realignment may dominate and is responsible for sensorimotor and cognitive after-effects (Michel, 2016; Michel, Pisella, et al., 2003). In force field adaptation however, realignment may be minor and responsible for short-lasting and poorly generalized sensorimotor after-effects (Cothros, Wong, & Gribble, 2006; Kawato, 1999; Kluzik, Diedrichsen, Shadmehr, & Bastian, 2008; Mattar & Ostry, 2007) and non-significant cognitive after-effects (Michel et al., 2018).

What is so peculiar to prism exposure and seems absent in force field adaptation? In the present investigation, we rolled back to a visuo-motor task which approaches the perturbations induced by prism exposure. The main difference between prism exposure and visuo-motor adaptation is the indirect inter-sensorial and sensorimotor conflict provided by the robotic device. We specifically controlled two important parameters. First, we induced either leftward or rightward 15-degree rotations. We adopted that angular amplitude to compare with other experimental protocols that use prism deviations (Michel, 2016). As for prism adaptation, we expected that adaptation to a leftward optical rotation would induce cognitive after-effects (Michel, 2016). Second, we presented the perturbation either abruptly (from one trial to the next) or gradually (over a certain number of trials). Critically, participants were aware of the perturbation in the abrupt condition but not in the gradual condition. Gradual exposure to the perturbation is known to strengthen the spatial realignment process (Jakobson & Goodale, 1989; Kagerer, Contreras-Vidal, & Stelmach, 1997; Michel, Pisella, Prablanc, Rode, & Rossetti, 2007). Further, representational after-effects depend on the realignment (Michel et al., 2003). We therefore reasoned that gradual exposure and not abrupt exposure to the perturbation would favor the occurrence of representational after-effects. According to this factorial design (direction of perturbation x type of perturbation), we hypothesized that representational after-effects would occur only following adaptation to a leftward rotation and would be more pronounced after gradual than abrupt exposure. Quite surprisingly, our results did not confirm our hypotheses. Instead, we found that none of the conditions produced significant cognitive after-effects. However, the sensitivity with which we discriminate the center of the line, that we called “representational acuity”, increased following the perturbation (irrespectively of the direction). This change was particularly marked after gradual exposure and persisted for some time after the sensorimotor adaptation.

## Materials and methods

### Participants

Fifty-six right-handed adults (26 women, 30 men, mean age = 23 years old, SD = 4.8) voluntarily participated in the experiment. All participants were healthy, without neuromuscular disease and with normal or corrected to normal vision. All participants gave their informed consent prior to their inclusion in the study, which was carried out in agreement with legal requirements and international norms (Declaration of Helsinki, 1964). All participants were naïve as to the purpose of the experiments and were debriefed after the experimental sessions.

### Experimental procedure and apparatus

The experiment consisted in three separate phases (BISECTION-PRE, ADAPTATION AND BISECTION-POST), described below. Briefly, we assessed space representation with a line bisection task before and after a visuomotor adaptation paradigm involving passing through targets with a haptic device. During the three phases, participants were comfortably seated in front of a virtual environment with the head on a chin rest in a dimly illuminated room. They looked into two mirrors that were mounted at 90 degrees to each other, such that they viewed one LCD screen with the left eye and one LCD screen with the right eye. This stereo display was calibrated such that the physical locations of the robotic arm was consistent with the visual disparity information.

During the BISECTION-PRE phase, participants were asked to perform a bisection segment test. A total of 130 horizontal green segments (length: 400 mm, thickness: 2 mm) were sequentially displayed on the screen together with a perpendicular red tick (height: 30 mm, thickness: 2 mm). Participants were requested to judge (forced-choice) whether pre-transected lines were bisected to the left or to the right of their center. The response was given verbally and manually recorded by the same experimenter. No response provided before a 20-second timeout was considered as missing information. Tick offsets were not uniformly distributed but followed a Gaussian shape in order to increase sampling granularity around the Euclidian center and, hence, enhance sensitivity of subsequent data analysis. To prevent any cognitive strategy, participants were told that ticks never occurred exactly on the middle of the segment and that their distribution was not systematically symmetric. Ticks were presented randomly and the computer screen went white for 1500 ms between each trial to reset the visual display and prevent the participant from using any cue between trials.

We ran a technical validation experiment to make sure that our bisection line test was sensitive to well-established stimuli. To do so, we enrolled an additional 13 participants (9 women, 4 men, mean age = 23.6 years old, SD = 4) who performed exactly the same test as the one used in our experiment. A total of 234 trials were presented in three blocks of 78 trials. In this case, however, a blue circle was displayed at the left or at the right of the segment in 33% of the trials, respectively. In the remaining third of the trials, no cue was visible. These three conditions varied randomly trial by trial. We found that the presence of the left cue significantly shifted the perceived center of the segment to the left compared to when there were no cues (paired t-test: t_12_=2.3, p=0.032, 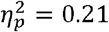). We found the same effect, but reversed, when the right cue was displayed (paired t-test: t_12_=2.5, p=0.022, 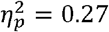). Our setup then reliably replicates the known cueing effects on space representation (e.g. Milner, Brechmann, & Pagliarini, 1992).

During the ADAPTATION phase, participants used the robotic arm to hit targets from the same starting position (white circle, diameter: 0.3 cm). One of five circular targets (diameter: 0.3 cm) appeared and were located on a circle centered on the starting position (radius: 20 cm) and were presented randomly vertically above the home position (0 deg), or at 10 and 20 degrees to the left (−10 deg and −20 deg) or at 10 and 20 degrees to the right (+10 deg or +20 deg). The 3D position of the robot handle was mapped in real time on a blue cursor (diameter: 0.2 cm). A trial started when the cursor was positioned inside the starting circle. We instructed and trained participants to reach the targets with a peak velocity within 65-75 cm/s. After each trial, the target turned red if peak velocity was larger than 75 cm/s or turned blue if peak velocity was smaller than 65 cm/s. The robot then gently pushed the hand back toward the starting position to prevent participants from planning and performing active movements.

Participants were randomly dispatched in four experimental groups of 14 participants according to the nature of the visuomotor perturbation (Fig. 1A). In the ABR+15 group, the perturbation consisted of a rightward 15-degree rotation of the cursor introduced abruptly (ABR), after 80 trials and for 280 trials. The ABR-15 group experienced the same condition except that the perturbation was opposite (−15 deg). The GRA+15 and GRA-15 groups performed first 76 trials. Then, the visuomotor rotation linearly increased (Fig. 1A, *Transition*) for 34 trials by steps of 0.429 deg to reach either +15 deg (GRA+15) or −15 deg (GRA-15). Then, participants underwent 250 trials under maximal rotation. All groups performed the same total number of trials (360). Participants in the ABR and GRA groups experienced cumulative visuo-motor perturbations of 4200 deg and 4005 deg respectively (in absolute value). The relative difference between the two groups is only 4.6%. Unlike in other adaptation paradigms, we decided to not expose participants to a washout phase in order to maintain their adapted state. When debriefed after completion of the full experimental session, participants were aware of the perturbation in the ABR condition but not in the GRA condition.

**Figure 1.**
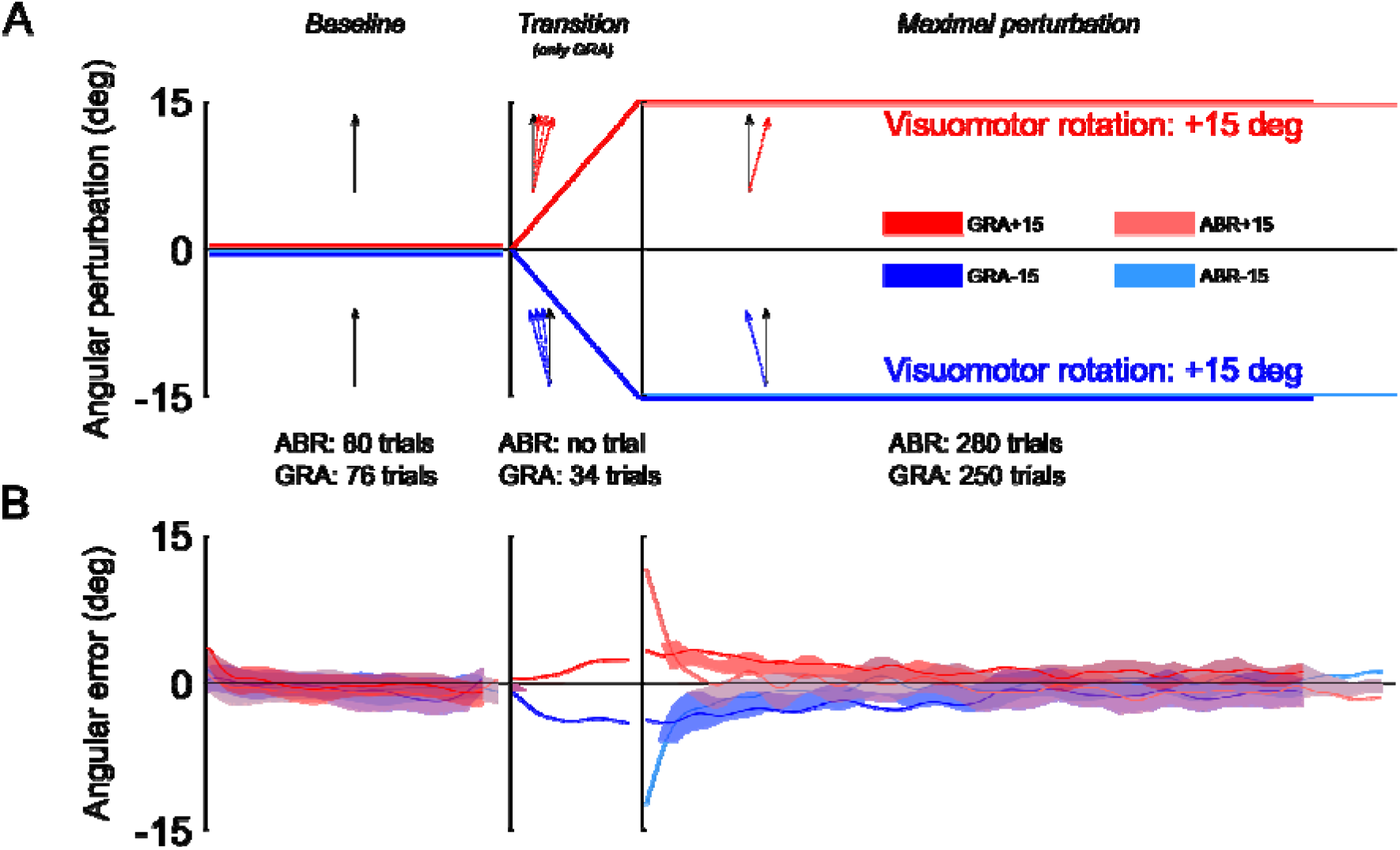
Illustration of the protocol and angular errors during the adaptation phase. (A) Participants in both groups were trained with pointing movements without visuomotor rotation (Baseline). During the transition, the GRA group was exposed to leftward or rightward visuomotor rotations that linearly ramped from 0deg to either +15deg or +15deg (transition) and then reached a maximal perturbation for the remaining trials (Maximal perturbation). After the baseline, participants from the ABR groups were immediately exposed to maximal perturbations, without transition. (B) Mean and standard error of angular errors measured during the baseline, the transition (for the GRA group only) and the maximal perturbation phases. Notice the high and mirror symmetric errors in the first trials of the Maximal perturbation phase for the ABR group and moderate but still mirror symmetric errors in the GRA group.

During the BISECTION-POST, the device, instructions and task were identical to those used in the BISECTION-PRE phase, except for the number of trials. Here, a total of 195 horizontal green segments were displayed on the screen and grouped in five blocks of 39 trials. We ensured again a balanced distribution of offsets within each block, therefore allowing us to assess a possible effect of the passage of time on space representation.

### Data analysis

In the BISECTION-PRE and BISECTION-POST phases, we recorded participants’ verbal responses for every trial (“Is the tick positioned to the right or to the left of the veridical segment midline?”). We then calculated the proportion of ‘RIGHT’ responses in function of the offset separately in each block (one single block in BISECTION-PRE and 5 blocks in BISECTION-POST). This S-shape function saturated at 0% for large negative (left) offsets and at 100% for large positive offsets (right).

To quantify the offset that corresponded to chance level (50%), i.e. the subjective perceptual estimation of the line center, we fitted logistic functions in each block, and for each participant separately (r^2^=0.91 on average), 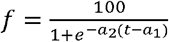, where a_1_ and a_2_ were regressed and t corresponds to the offset. The subjective line center that corresponds to f=50 is a_1_.

Restricting the analysis of the distributions of the responses only to the offset is too limitative. Indeed, different logistic regression functions can yield the same subjective line center. Figure 2 depicts three simulated examples of logistic regression lines (f, g and h). One can see that functions f and h intersect at exactly the ordinate of 50%, and therefore lead to the same offset (−3 cm). Function g has a different offset (4 cm). We therefore also quantified the sensitivity of the decision curve with the derivative of the model. We define the perceived representational acuity as Δ*w*. This parameter is calculated as the difference between two values of w, defined as 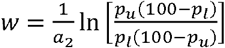, with [*p*_l_;*P*_u_] = [40%; 60%] and Δ*w* = *w*_u_ — *W*_l_. For a given probability of response (20% centred on 50%, or chance level), a small Δ*w* means that the regression curve is steep and that one can decide with high sensitivity (certainty) whether an offset is to the right or to the left of the center. In contrast, a large Δ*w* reflects a flatter regression curve and a low sensitivity (uncertainty) to discriminate between left and right ticks. An extreme (fictious) case would be represented by a horizontal flat regression line and would mean that the change to identify the positions of segments is purely dictated by chance (in that case, Δ*w* = +∞). The smaller Δ*w*, the better the perceived representational acuity.

**Figure 2.**
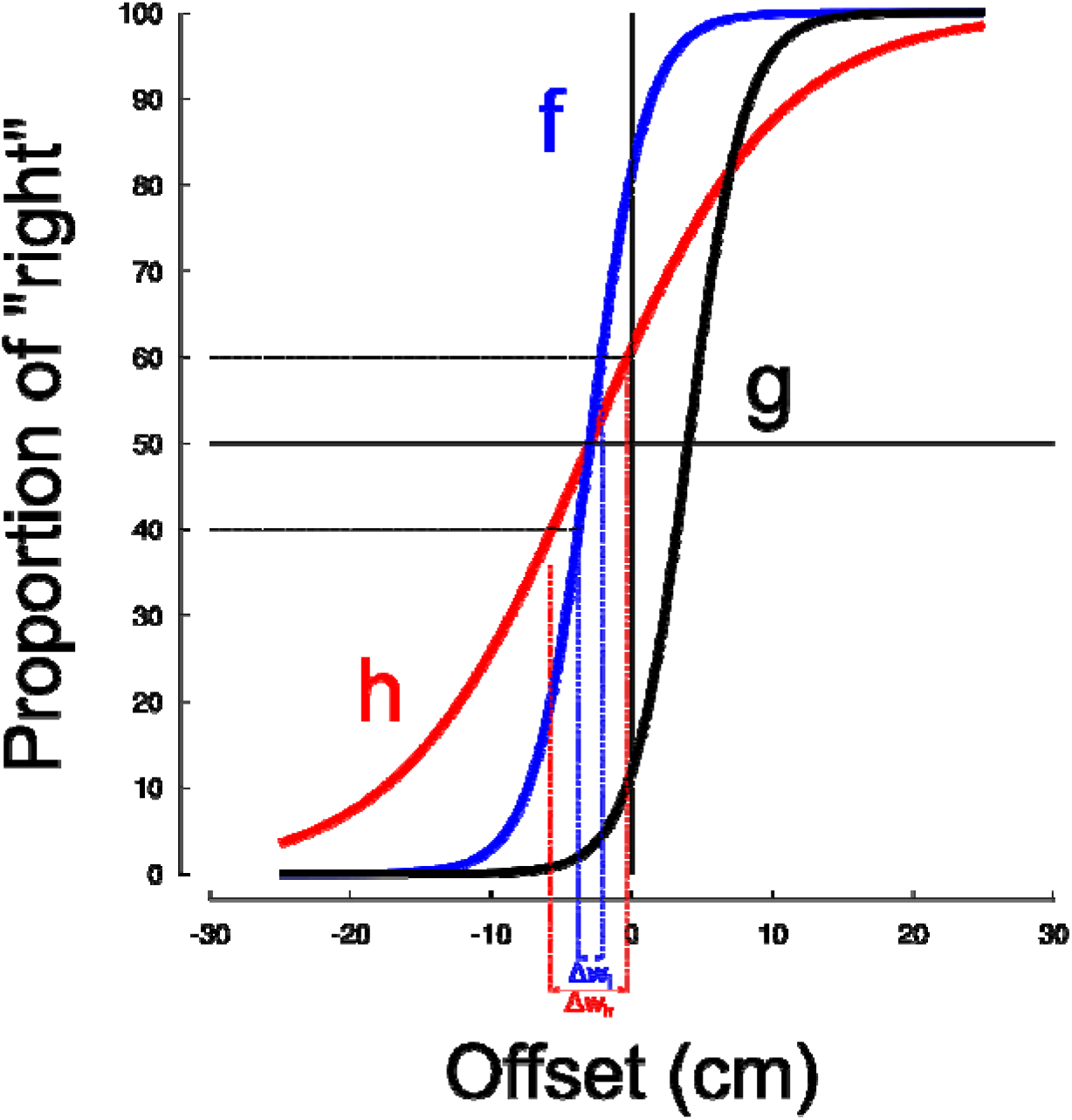
Simulation of logistic functions. Functions f and g are parallel and yield different offsets. Functions g provides the same offset as function h but is different. Functions were defined as — — —.

During the ADAPTATION phase, cursor positions were recorded with a sampling rate of 500 Hz. Movement start was detected when movement velocity exceeded 3 cm/s for at least 100 ms. Direction error of each movement was defined as the angle between straight ahead and the segment connecting the start position to the position of the cursor 150 ms after movement onset.

Quantile-quantile plots were used to assess normality of the data before using parametric statistical tests. Variables of interest were submitted to different statistical models (repeated measures ANOVAs) according to the effects analyzed. Independent t-tests were conducted to compare data between groups and paired t-tests were used to compare different conditions within groups. We also report partial eta-squared for significant results to account for effect size. Data processing and statistical analyses were done using custom routines in Matlab (The Mathworks, Chicago, IL).

## Results

We designed an experiment to test if the transfer of sensorimotor effects induced by visual rotation to spatial representation is a phenomenon peculiar to prism adaptation. Participants were confronted to either leftward (L) or rightward (R) visuomotor rotations during reaching movements. The perturbation was introduced abruptly (ABR) or gradually (GRA). Different participants were attributed to four groups following a factorial design (G_L,ABR,_ G_L,GRA,_ G_R,ABR_ or G_R,GRA_). Participants performed a line bisection test before and after the adaptation phase.

### Groups adapted to gradual and abrupt visual motor rotations

Each group experienced a baseline sequence during which 80 trials were performed to one of five targets (Fig. 1B, “Baseline”). No rotations were introduced in that sequence. The errors were on average −0.3deg (SD=1.4deg). Within each group, the mean error was not different from 0deg (independent t-test, G_L,ABR_: t26=0.06, p=0.950; G_L,GRA_: t_26_=-1.89, p=0.071, G_R,ABR_: t_26_=-1.20, p=0.243 and G_R,GRA_: 1036, p=0.187) and group performances were comparable in terms of direction errors (all t<0.28, all p>0.272).

Two groups were exposed to abrupt perturbations (G_L,ABR_ and G_R,ABR_) and two groups experienced gradual perturbations (G_L,GRA_ and G_R,GRA_). Participants in the two ABR groups (left and right perturbations) exhibited large errors during the first trials in the “Maximal perturbation” sequence (Fig. 1B, light colors). These errors reached −12.9deg in G_L,ABR_ and 13.5deg in the G_R,ABR_. They reached the same absolute amplitude (t_26_=-0.65, p=0.520) and were different from 0 (G_L,ABR_: t_26_= 16.80, p<0.001, η^2^=0.92 and G_R,ABR_: t_2_6=21.75, p<0.001, η^2^=0.95). Participants in the GRA group also showed errors in the early trials of the “Maximal perturbation” sequence (Fig. 1B, dark colors; G_L,GRA_: −4.8deg and G_R,GRA_: 3.7deg). However, their amplitudes, although the same in absolute values (t_26_=0.78, p=0.443), were smaller than in the ABR groups (t54=10.61, p<0.001, η^2^=0.68) and also different from 0deg (G_L,GRA_: t_26_=4.01, p<0.001, η^2^=0.39 and G_R,GRA_: t_26_=5.06, p<0.001, η^2^=0.51). In contrast to the ABR groups, the GRA groups were exposed to a 34-trial transition during which the rotation of the cursor linearly went from 0deg to −15deg for G_L,GRA_ or +15deg for G_R,GRA_ (Fig. 1B, “Transition”). Unlike in the ABR groups, participants of the GRA groups did not report having detected any sensorimotor disturbance while using the robot. Statistics did not report significant differences between the last trials in the “Transition” sequence and the errors in the first trials of the “Maximal perturbation” sequence (G_L,GRA_: t_13_=0.91, p=0.381, η^2^=0.39 and G_R,GRA_: t_13_=1.33, p=0.206).

At the end of the long “Maximal perturbation” sequence, all groups were adapted to the visual rotation. Indeed, the errors during the last 3 trials were on average 0.03deg (SD=2.54deg) and were not significantly different from 0deg (all t<1.9, all p>0.710).

### Effects of sensorimotor adaptation on space representation

We quantified the effect of visuomotor adaptation on space representation by means of a standard line bisection task. Figure 3 depicts subjective line center values regressed through the sigmoid model before (“Pre”, light colors) and after (“Post”, dark colors) sensorimotor adaptation separately in the ABR groups and GRA groups. In addition, the series are presented separately for the left (−15, blue) and right (+15, red) perturbation conditions. Together, the four groups had initial rightward (0.16) estimation of line center (all different from 0, t_94_=4.7, p<0.001, η^2^=0.19). This analysis held true at the group level (all t>2.1, p<0.045) except for the ABR+15 group (t_22_=1.9, p=0.076, η^2^=0.14). All groups were however comparable (all t<0.4, p>0.656) before entering the visuomotor adaptation task.

**Figure 3.**
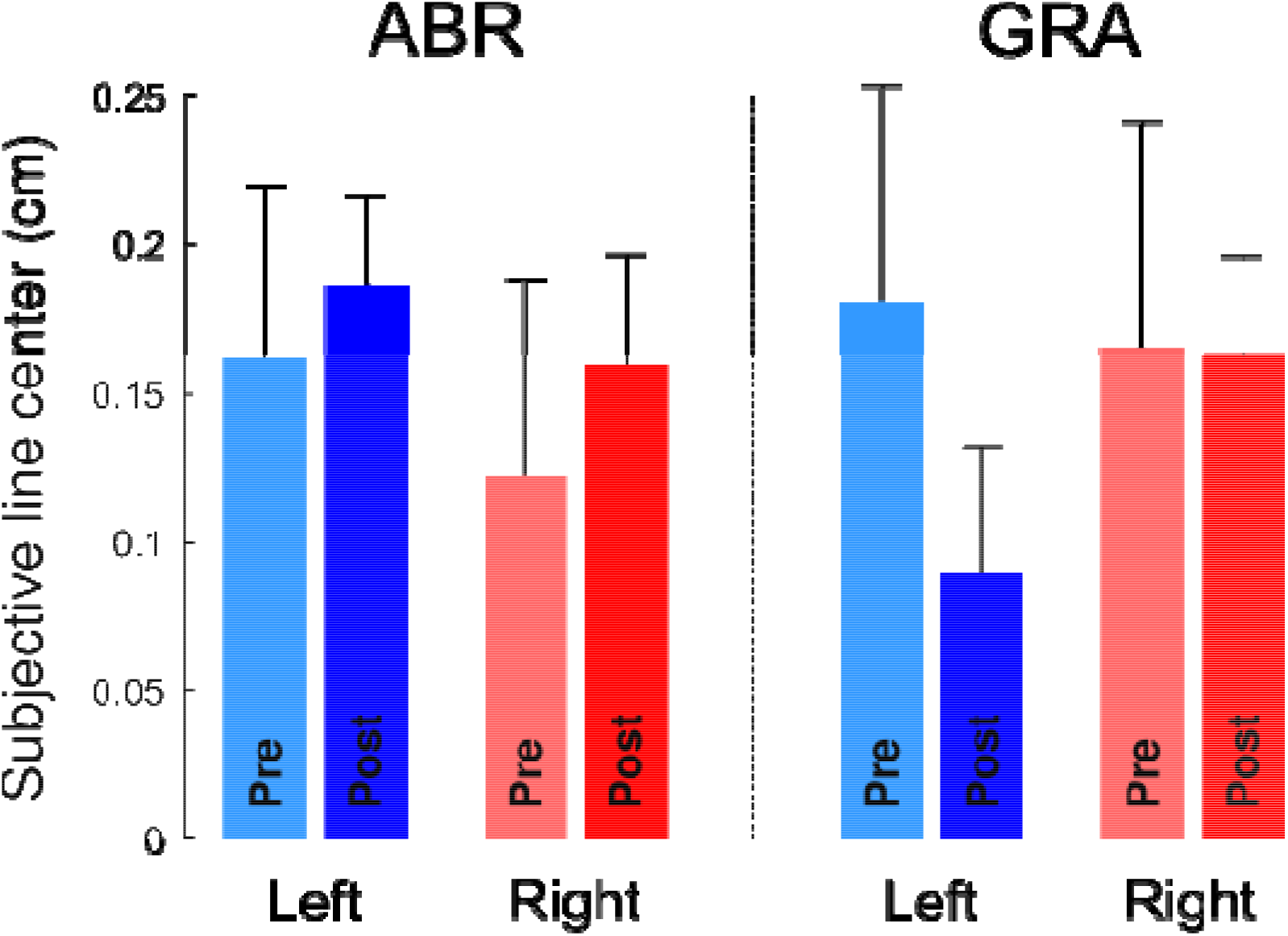
Subjective line center in the group exposed to abrupt (ABR) and gradual (GRA) perturbations. Blue colors correspond to leftward cursor rotations (−15deg) and red colors correspond to rightward cursor rotations (+15deg). Subjective line center before and after the adaptation phase are denoted by light and dark colors respectively. No conditions affected this value.

We made a statistical analysis to quantify the effects observed in Figure 3. We ran a RM-ANOVA with the factors modality (ABR vs. GRA), direction of perturbation (Left vs. Right) and Time (Pre vs. Post). None of these factors influenced space representation (all F<0.5, all p>0.474) and no interaction was significant either (all F<1.3, all p>0.255).

We attempted to highlight any difference that would emerge with time within the Post period. However, again, we failed to highlight such effect of the passage of time on subjective line centers across the five post blocs. This was quantified by a 1-way RM ANOVA with factor post bloc (1 to 5) and yielded F_4,163_=0.7, p=0.608.

Any measurement scale is characterized by its graduations but also by its sensitivity to the underlying phenomenon. In this context, while our ability to perceive the position of a tick on a segment may not be influenced by any of our experimental conditions, the sensitivity of the perceptual system itself may well be altered. In other words, experimental conditions could blur or improve the clarity of our ability to discern midline. To assess this, we tested how representational acuity (Δw, see Methods) was influenced by experimental conditions. Figure 4 depicts Δ*w* in function of direction, modality and time. The RM ANOVA with factors Direction, Modality and Time reported a main effect of modality (F_1,111_=7.1, p=0.009, η^2^=0.05) and time (F_4,111_=4.9, p=0.0004, η^2^=0.16) on Δ*w* but not direction (F_1,111_=2.1, p=0.155). Post hoc tests showed that Δ*w* dropped (representational acuity improved) after the adaptation phase (pairwise comparisons; all t>2.54, all p<0.014) and remained constant throughout the 5 post blocs (pairwise comparisons of the five post blocs; all t<0.5, all p>0.619). There was no interaction between factors (all F<1.74, all p>0.189) and the four groups had comparable Δ*w* before adaptation (all t<1.162, all p>0.261). Altogether, this analysis shows that representational acuity improved after the adaptation phase and was even slightly better after gradual exposure. Furthermore, representational acuity remained stable with time in the post-adaptation blocs.

**Figure 4.**
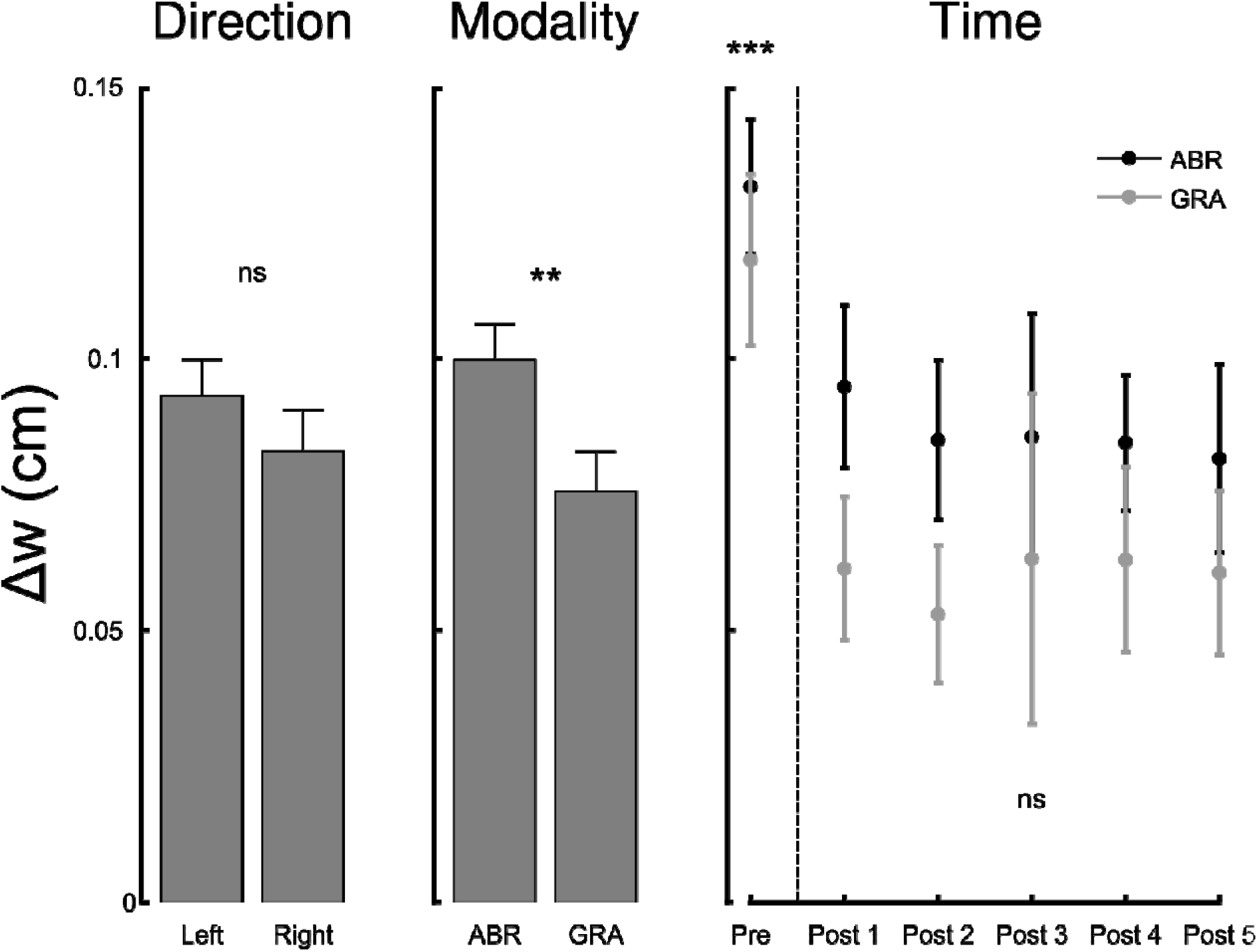
The width of the interval in which participants provide reliable responses to the line bisection task varies with modality (ABR, Abrupt and GRA, Gradual) and time but not direction of perturbation (L, Left vs. R, Right). The narrower the interval, the best the representational acuity. The x-axis of the right panel (Time) corresponds to the pre-adaptation block and the five post-adaptation blocs. Representational acuity improves (Δw decreases) after exposure to visuo-motor rotations and remains stable after. ns=not significant, **=p<0.01, ***=p<0.001.

## Discussion

The main objective of the present experiment was to investigate whether adaptation to visuo-motor rotation transfers to space representation in healthy individuals. Despite optimal conditions to develop sensorimotor adaptation and sensitive conditions to investigate spatial representations, we failed to observe any representational after-effects. Nevertheless, we pointed out that the sensitivity with which we discriminate the center of the line is improved following the perturbation.

The identification of processes that encompass sensorimotor plasticity could help to understand the favorable conditions to produce cognitive after-effects. As reviewed in the introduction, prism adaptation is responsible for after-effects on space representation in healthy individuals (see Michel, 2016 for a review) whereas adaptation to visuo-motor rotation used in the present study is unable to produce such representational after-effects. The occurrence of cognitive after-effects depends on the presence of sensorimotor after-effects (Michel, Pisella, et al., 2003). Therefore, the understanding of the conditions favorable to developing sensorimotor after-effects and their generalization beyond the perturbation apparatus to a broader spatial context could help us to understand the transfer of sensorimotor plasticity to spatial cognition. A recent and inescapable review of the literature comparing the sensorimotor processes with respect to specific methodological features provide useful understanding keys (Fleury, Prablanc, & Priot, 2019).

One first has to consider the relative contribution of adaptive processes in behavioral changes. When a motor commanded is issued, the forward model receives a copy of this motor command (efference copy) and makes predictions about the sensory consequences of the upcoming action. The discrepancy between actual and predicted reafferences (prediction error) acts as an error signal that triggers adaptive processes to maintain accuracy of the controller and the forward models (Kawato, 1999; Wolpert & Miall, 1996). During early exposure to the perturbation, when the system is subject to large errors, the rapid calibration process occurs. It improves the accuracy of the action by processing the error signal. In parallel, the slow realignment process brings the sensory and motor reference frames back in congruence. Calibration is cognitively demanding and contributes little to after-effects. In contrast, realignment is more automatic and develops gradually. Realignment is thought to be mainly responsible for after-effects. Because the occurrence of cognitive after-effects depends on the development of the realignment (Michel, Pisella, et al., 2003), we may hypothesize that a fundamental difference between prism and visuo-motor adaptations may lie in the relative contribution of the recalibration and realignment processes. The recalibration process may dominate in adaptation to visuo-motor rotation, entail poorly generalized sensorimotor after-effects (Ghahramani, Wolpert, & Jordan, 1996; Pine, Krakauer, Gordon, & Ghez, 1996) and develop non-significant cognitive after-effects. We used a protocol based on gradual exposure to visuomotor rotations to maximize the chances to induce spatial realignment (Jakobson & Goodale, 1989; Kagerer et al., 1997; Michel, Pisella, et al., 2007). However, these otherwise favorable experimental conditions appeared to be insufficient to produce representational after-effects.

Second, we have also to consider the context in which adaptive processes develop. Although exposure to prisms and visuo-motor rotations imply comparable visual mismatches between expected and actual reafferences, they differ by the nature of the reafferences. During prisms exposure, reafferences are direct and real without interface and actions are body-centered. The lack of interface involves adjustments that are context-independent. On the contrary, during visuo-motor rotation exposure, reafferences are symbolic and indirect through a robotic arm and a computer interface. In that case, actions are not body-centered. Whether or not participants hold a (passive) robotic arm allows them to switch control (Kluzik et al., 2008). Adaptation to visuo-motor rotation may rely on a context-dependent process because this adjustment would be specific to the interface (Fleury et al., 2019). The concept of generalization corresponds to the persistent compensations that can be observed in different contexts from exposure conditions (Bastian, 2008; Censor, 2013; Poggio & Bizzi, 2004). It depends considerably on specific exposure conditions. Robust generalization of after-effects outside the exposure conditions reflects an adaptive process which is context-independent following prisms exposure (Bedford, 1989; Fleury et al., 2019). Conversely, the limited generalization to other conditions is associated with a high contribution of contextdependent processes following visuomotor rotation. A narrow spatial tuning is observed, with a sharp gradient around the trained direction highlighting a poor generalization across multiple directions (Ghahramani et al., 1996; Pine et al., 1996). This could be the reason why generalization did not occur in a broader spatial, cross modal and even representational context in line bisection task.

The lack of representational after-effects following visuo-motor rotation is the first important result of the present study that has strong methodological implications. Indeed, similar to adaptation to force field (Michel et al., 2019), adaptation to visuo-motor rotation reveals to be an appropriate tool to investigate specifically sensorimotor plasticity without producing any change in space representation assessed with line bisection. Therefore, visuo-motor rotation will be particularly appropriate to study sensorimotor plasticity in individuals who exhibit an inherent bias in space representation (e.g., hyperpseudoneglect in patient with schizophrenia or inverse pseudoneglect in children with dyslexia) (Michel, Bidot, Bonnetblanc, & Quercia, 2011; Michel, Cavezian, et al., 2007) while preserving it from any aggravation of their representational bias.

Another strong and innovative result of the present study was the improvement of the sensitivity with which participants visually discriminate the center of the lines, that we proposed to call ‘representational acuity’. Indeed, participants’ representational acuity improved after the adaptation phase and was even slightly better after gradual exposure. The representational acuity was also previously considered as the indifference zone, i.e. the portion of line that is considered as the center of the line (Manning, Halligan, & Marshall, 1990; Nicholls & Roberts, 2002). It has been shown that this acuity depends on the configuration of the stimulus/line to bisect (Westheimer, Crist, Gorski, & Gilbert, 2001). The notion of acuity has also been investigated in somatosensory perception in participants who made active reaching movements or matched passive movements to an unseen target using a robot arm. When perceptual testing procedure targeted proprioceptive function, somatosensory experience paired with reinforcement improves perceptual acuity to a degree comparable with that seen following training with active movements (Bernardi, Darainy, & Ostry, 2015). In our experiment, the visual modality was concerned. Representational acuity was improved following exposure to a visuo-motor rotation meaning that sensorimotor adaptive/learning processes improve visual representational acuity. This improvement also persists for some time; we did not find any reversing trend of this variable. Our results underline the strong link between sensorimotor plasticity and representational acuity. It is then likely that when there is an intersensory conflict (even indirect in the case of visuo-motor rotation), the sensory modalities exposed and involved in the recalibration / realignment processes would become more sensitive. Thus, the ability to discriminate the center of a line would be exacerbated. Furthermore, in a general view, our results highlight also the dissociation between the after-effects in space representation (the lack of the displacement of the subjective line center) and after-effects in representational acuity (the increase of the sensitivity with which the participants discriminate the center of the lines) following visuo-motor rotation exposure.

## Conclusion

We have shown two main results with strong theoretical and methodological consequences. First, in the specific conditions that we used, visuo-motor rotation could be considered as an appropriate tool to investigate sensorimotor plasticity while sparing representation level. Second, exposure to visuo-motor rotation could be considered as a useful tool to increase the representational acuity. Follow-up experiments testing participants under different experimental procedures will allow us to better define the sensorimotor conditions necessary to modulate the spatial representation and to increase representational acuity.

## Acknowledgments

This research was supported by the « Institut National de la Santé et de la Recherche Médicale » (INSERM), the « Conseil Général de Bourgogne » (France) and the « Fonds Européen de Développement Régional » (FEDER). The authors wish to thank J. Ferrera for her efficient assistance to collect data for the technical validation of the setup.

